# Systematic chemical genomic screening of *Vibrio cholerae* maps gene-phenotype relationships

**DOI:** 10.64898/2026.01.16.699870

**Authors:** James RJ Haycocks, Huda Ahmad, Micheal Alao, Georgia Williams, Susan Sutherland, Chris LB Graham, Laura Alvarez, Prateek Sharma, Patrick J Moynihan, Felipe Cava, David C Grainger, Danesh Moradigaravand, Manuel Banzhaf

**Author notes:** Contributed equally (James RJ Haycocks, Huda Ahmad, Micheal Alao, Georgia Williams). DATA AVAILABILITY: The chemical genomics dataset is available in a browsable format from https://chemgenxplore.kaust.edu.sa/.

## Abstract

*Vibrio cholerae*, the causative agent of cholera, remains a major global health burden despite ongoing initiatives for vaccination and improved sanitation. Much of the *V. cholerae* life cycle occurs in aquatic environments, requiring gene functions that enable survival under diverse stresses associated with this niche and during the transition to the human host. Although numerous *V. cholerae* genome sequences are available, functional annotation across its pan-genome remains incomplete, and many annotated genes lack experimental validation.

In this study, we employed a chemical genomics approach to profile fitness phenotypes across 104 diverse stress conditions using a library of 3,026 single-gene mutants of *V. cholerae* C6706, a widely used research strain. In total, we identified significant growth phenotypes for 1,518 mutants, of which 88 correspond to unannotated (hypothetical or putative) genes. Together, these data provide a comprehensive resource for exploring gene function in *V. cholerae*, supported by detailed quality metrics and open-access tools to facilitate community-driven discovery.

**AUTHOR SUMMARY:** Although many bacterial pathogens have been extensively sequenced, we still know relatively little about what many of their genes actually do. In many cases, gene functions are solely predicted by computer algorithms but have not been confirmed in the laboratory, and we can assume that some of those predictions may be incorrect. Chemical genomics — a method that tests thousands of single-gene mutants under different stresses — can help reveal what these genes are responsible for. Using this approach, we screened a library of *Vibrio cholerae* C6706 mutants under 104 different conditions. We found that more than half of the mutants showed measurable effects on growth, including many genes that were previously uncharacterized. Here, we describe our experimental pipeline, present key quality checks, and share how other researchers can use this dataset as a resource to explore *V. cholerae* mutant phenotypes.

## INTRODUCTION

*Vibrio cholerae* is a Gram-negative bacterium responsible for causing the disease cholera, of which there are estimated to be 4 million cases each year, resulting in 140,000 deaths (1). Global case numbers are rising, and the disease is endemic in many low-income regions (particularly in parts of Africa and Asia), or in settings where sanitation systems have failed (2,3). A key challenge in managing the disease is the biphasic life cycle of *V. cholerae*, which has a large aquatic reservoir in coastal and estuarine habitats (4–6). Here, the organism forms biofilms on chitinous surfaces, undergoes natural transformation, and readily assimilates new genetic material (7–9). Upon entering the human host, *V. cholerae* initiates a virulence programme that enables colonisation of the small intestinal crypts (10,11). The production of virulence factors such as cholera toxin (CT) results in subsequent chloride ion and water loss into the intestine, causing profuse watery diarrhoea (12,13).

Thousands of *V. cholerae* genome assemblies are currently available from the NCBI database, including many for strains belonging to the El Tor biotype, which is responsible for causing the seventh and current cholera pandemic (14). However, functional annotation of this pan-genome lags behind sequencing efforts. Uncharacterised genes are often assigned functions computationally rather than experimentally. Although such homology-based annotation is helpful for prioritising candidate genes, it remains error-prone (15–17), and experimental validation is required for definitive assignment. For example, in the original genome annotation of the first-sequenced strain of *V. cholerae*, N16961 (El Tor biotype), 40.6% of open reading frames were classified as hypothetical, thus lacking any functional annotation (18). Whilst this proportion has been reduced to 6.2% in a recent update through the NCBI Prokaryotic Annotation Pipeline (19), a substantial number of genes still lack experimental validation, which hinders our understanding of *V. cholerae* biology and, in turn, downstream efforts to understand and combat cholera.

Chemical genomics systematically profiles the growth of arrayed single-gene mutant libraries under defined chemical and environmental conditions to infer gene function (20). This approach complements homology-based annotation by linking individual mutant phenotypes to functional networks, enabling the mapping of stress responses and metabolic pathways and identifying genes important for fitness under specific conditions. Chemical genomics approaches have been especially effective in model organisms such as *Escherichia coli* K-12, *Bacillus subtilis* and *Saccharomyces cerevisiae* (21–25). In bacterial pathogens, chemical genomics screens have typically used pooled mutant libraries to probe gene-drug interactions (26–28). In *V. cholerae*, arrayed single-mutant libraries have been previously used for targeted screens, including studies identifying motility genes and small-molecule inhibitors of the virulence regulator ToxT (29,30).

In this study, we performed a chemical genomics screen using an ordered single-gene transposon insertion mutant library of the widely used El Tor strain *V. cholerae* C6706 (31). Across 104 conditions, we identified at least one significant fitness phenotype for 1,518 of the 2,748 gene mutants screened (55.2%). Of these, 1,430 mutants (94.2%) corresponded to annotated genes, and 88 mutants corresponded to unannotated genes (5.8%). We present quality control and key screening metrics, and demonstrate how this dataset can be mined to assign putative functions to hypothetical genes. We also illustrate how the scientific community can browse and explore the data to prioritise their own follow-up studies through ChemGenXplore, a Shiny app with a graphical user interface that enables interactive access and visualisation (32).

## METHODS

### Mutant library re-annotation

For this study, we used an ordered single-gene transposon insertion mutant library of *Vibrio cholerae* generated by Cameron *et al.* (31). The library was created by introducing TnFGL3 into a C6706 Δ*lacZ* parent strain of the El Tor biotype, which was originally isolated from Peru in 1961 (33). Although insertion sites for this library were originally determined using the original annotation for the *V. cholerae* N16961 genome, they were remapped here to a more recent re-annotation of this genome (accession number GCF_000006745.1, PGAP annotation RS_2025_08_23). Information on clone identity, well position, and transposon insertion sites was obtained from Cameron *et al.* (31), and remapping was performed using basic R scripts to merge and cross-reference genomic coordinates and annotation tables, thereby assigning insertion sites to the corresponding updated loci and genomic features. The resulting dataset was further cross-referenced with UniProt (34) and eggNOG (35) databases to retrieve predicted protein function annotation.

Finally, the dataset was manually curated to ensure consistency between genomic coordinates, locus assignments, and functional annotations prior to integration into the revised dataset and downstream analyses.

### Library condensation

Prior to the screen, the library was condensed to 3,026 mutants with consistent visible growth in LB broth, and mutants exhibiting inconsistent or no growth in LB were excluded from the analysis. A list of all mutants used is provided in Table S1. The screening library used also included 21 replicates of wild type *V. cholerae* C6706.

Using an Opentrons OT-2 liquid handling robot (Opentrons Labworks Inc.), 10 µl of each culture from the original 96-well plates was transferred into new 96-well plates containing 100 µl LB broth per well. Plates were sealed with breathable seals and incubated for 14 hr at 37°C and 180 rpm. Subsequently, 40 µl of 50% glycerol was added to each well, and the plates were sealed with aluminium film and stored at-80°C. For library condensation, LB (Lennox) broth (Sigma-Aldrich) was used. To prepare solid media, 20 g/l agar (Sigma-Aldrich) was added.

### Chemical genomics screen

We used a chemical genomics screening pipeline as previously described (20). Briefly, the ordered library was arrayed in high-density (1,536 mutants/plate) on 2% LB Lennox agar plates (Sigma-Aldrich) using a BM3-BC robot (S&P Robotic Inc.). Plates were incubated at 30°C for 8 hr. These plates, referred to as source plates, were subsequently used to pin the arrayed library onto condition plates containing the stressor.

A full list of all conditions used and their categorisation is provided in Table S2. Condition plates consisted of 2% LB Lennox agar, except for nutrient starvation conditions, for which M9 minimal medium (Sigma-Aldrich) was used. Chemical stressors were mixed with agar during plate preparation. Environmental stresses were induced during incubation. Congo red (20 µg/ml) and Coomassie blue (10 µg/ml) dyes were added to all agar plates to enhance visibility of bacterial colonies during imaging (36).

Plates were incubated at 30°C for 8–10 hr until distinct colonies were visible. Plate imaging was carried out under controlled lighting using an 18-megapixel Canon RebelT3i (Canon) camera mounted on the BM3-BC robot (S&P Robotic Inc.). The resulting images were analysed with Iris software (36), using endpoint colony size as the readout. Iris automatically identifies plate edges, segments colonies, measures colony pixel areas, and assigns numeric values for the size of each colony. Each condition was tested using at least three biological replicates.

### Data analysis and quality control

Iris data were processed using ChemGAPP software (37), which applies several quality control checks, including curation by removal of poor-quality conditions or other error such as mis-pinning or mis-labelling. ChemGAPP first carries out normalisation for edge effects using a Wilcoxon rank-sum test (38) to correct for colonies at the plate edges, which tend to be larger due to reduced competition (39). In addition, the software normalises colony sizes by scaling to the median colony size at each plate centre, producing values that are comparable across different conditions. Finally, ChemGAPP assigns fitness values (S-scores) (40) to the data based on colony fitness, by comparing the fitness of a mutant in each condition to the median fitness of the mutant in all screening conditions. This enables meaningful comparison of mutant fitness in different conditions whilst controlling for changes in baseline fitness.

Following this, several quality control tests are applied to identify unreliable data points and guide data curation. Briefly, the first of these uses a Z-score analysis to detect whether colonies have been mis-pinned or misidentified by Iris by detecting plate replicates that are outliers (or are missing) from the expected distribution of standard deviations. Secondly, a Mann-Whitney test (41) is used to identify plates that have been mislabelled, or unevenly pinned, by comparing colony sizes between replicate plates. As a result, mislabelled replicate plates will not fit the expected distribution. Finally, comparison of variance for Mann-Whitney p-values between replicate plates, and variance analysis between replicate colonies for each condition are used to identify non-reproducible conditions, which are removed from subsequent analyses.

To assess replicate reproducibility, pairwise Pearson correlations (42) of colony size measurements were calculated between all replicates, with higher correlation values indicating stronger agreement and values around r ≥ 0.6–0.8 commonly interpreted as strong reproducibility (43,44). The false discovery rate (FDR) was controlled using the Benjamini-Hochberg procedure (45) at 5%, hence phenotypes with an FDR of ≤0.05 were deemed to be significant.

### Phenotype count and cumulative distribution analysis

Gene-condition phenotypic data were obtained from S-scores, with statistical significance assessed using false discovery rate (FDR) correction. Significant phenotypes were defined as gene-condition pairs with an FDR ≤ 0.05. For each gene, a phenotype count was calculated as the number of unique conditions in which the gene exhibited a significant phenotype. Cumulative distributions were computed by calculating, for each phenotype count, the fraction of genes with phenotype counts greater than or equal to that count. All analyses were performed in R.

### Phenotypic correlation analysis

Gene-gene phenotypic correlations were calculated using Pearson correlation coefficients (42) across all screened conditions, retaining positive correlations with r > 0.4 to focus on moderate-to-strong associations and reduce noise. Gene pairs were classified as unannotated–annotated or annotated–annotated. Correlation coefficients were binned by correlation strength, and for each bin the fraction of gene pairs was calculated relative to the total number of pairs within each class. Pearson correlation coefficients (42) and associated p-values were computed per gene pair; p-values were summarised per correlation bin (using the median) for display purposes only. All analyses were performed in R.

### Conditionally essential genes

Conditionally essential genes were defined as genes whose mutants exhibited strong negative fitness effects under at least one tested condition, as indicated by highly significant negative S-scores (S-score ≤-10, FDR ≤ 0.05). This threshold was empirically supported by visual inspection of raw colony images, where mutants with S-scores ≤-10 consistently showed severe growth defects or absence of colony formation.

### Liquid growth curve assays

To assess the fitness of strains in liquid culture, growth curve assays was performed. Single colonies of *V. cholerae* from the single-gene mutant library were grown overnight in 5 ml LB medium, and the optical density at 600 nm (OD_600_) was measured. Cultures were then diluted to an OD_600_ of 0.05 using fresh LB medium. For stress condition assays, 5 µl of the selected stress agent was added to 5 ml of LB medium (concentrations are detailed in the figure legends).

Each experiment included multiple controls: a wild type strain without stress, a mutant knockout strain without stress and a control containing LB medium only. Test conditions consisted of the wild type strain exposed to the stress condition and the mutant knockout strain exposed to the same stress condition. For each condition, 200 µl of prepared culture was added to individual wells of a 96-well microplate, and all conditions were run in triplicate to ensure reproducibility. The plate reader was set to 37°C with double orbital shaking at 200 rpm. OD_600_ readings were recorded every 30 minutes over a 12 hr period, resulting in a total of 25 measurement cycles.

### Network analysis

A gene–gene network was constructed by computing pairwise Pearson correlations (42) between phenotypic profiles across all screened conditions. Gene pairs exhibiting strong phenotypic similarity (r ≥ 0.6) (43,44) were retained as edges in the network. Louvain community detection identified three dominant clusters of phenotypically correlated genes, with unannotated genes embedded within annotated gene communities (46). All network construction, analysis, and visualisation were performed in R using the igraph package (v2.2.1) (47).

## RESULTS

### Chemical genomic screening pipeline and screening conditions

Across sequenced *Vibrio cholerae* O1 El Tor genomes, including N16961 and the closely-related strain C6706, approximately 6% of genes are hypothetical genes that lack annotation (18,48) (N16961: GCF_000006745.1, PGAP RS_2025_08_23; C6706: GCF_009763945.1, PGAP RS_2025_12_20). In addition, many annotated genes are often assigned functions based solely on homology and lack experimental validation. A central aim of this study is to strengthen functional annotation by using chemical genomics to identify phenotypes for single-gene mutants exposed to diverse stressors. To achieve this, we used an ordered mutant library previously generated by Cameron *et al.* (31). The positions of transposon insertions in this library were mapped to the original annotation of the N16961 genome that was available at the time of construction. To update this, we subsequently re-mapped insertion sites to the revised annotation and manually curated the resulting locus assignments.

To carry out the chemical genomics screen, we used a high-throughput screening pipeline summarised in Figure 1A. Single-gene transposon insertion mutants from the *V. cholerae* C6706 library were first arrayed at high density (1,536 colonies per plate) to generate source plates.

**Figure 1.**
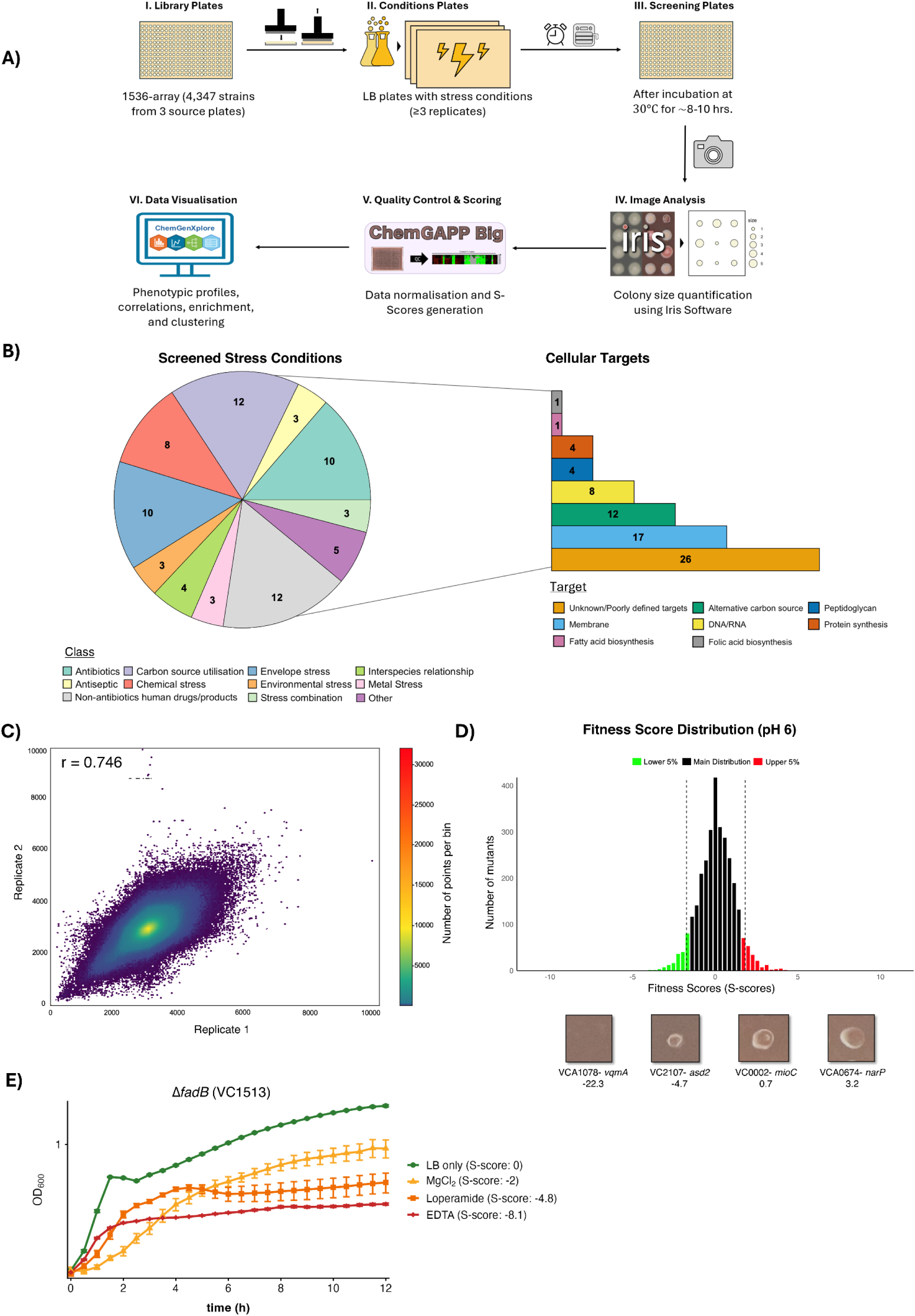
Chemical genomics screening pipeline and quality control of *V. cholerae* C6706 dataset. A) Summary of screening pipeline; I) The single-gene mutant library was arrayed in high density (1536 colonies/plate) onto source plates. II) Condition plates were made containing the stressor, and mutants were pinned onto these from source plates. III) Condition plates were incubated at 30°C (environmental stressors were applied during incubation if required) and subsequently imaged. IV) colony sizes were assigned numeric values by Iris, V) ChemGAPP Big software was used to calculate fitness (S-scores) for each mutant in each condition. VI) Downstream data analysis (such as phenotypic clustering) and visualization can subsequently be carried out using software such as ChemGenXplore. B) Pie chart showing classification of screened stress conditions (left) and their cellular targets (right). Note that some conditions have multiple cellular targets. C): Heat-map scatter plot showing size correlation between replicates. D) Histogram showing distribution of S-scores across all mutants in pH 6. The lowest 5% of S-scores (indicating reduced fitness) are shown in green, the top highest 5% of S-scores are shown in red (indicating increased fitness). Sub-panels below the graph show example images of mutant colonies with S-scores displayed below. E) Growth curves of *V. cholerae* C6706Δ*fadB* (VC1513) in liquid media, using conditions exhibiting different S-scores. Bacteria were grown in microtiter plates at 37°C, in LB media containing no additive (green circles), 5 mg/ml MgCl_2_ (S-score:-1.95, yellow triangles), 60 μg/ml loperamide hydrochloride (S-score:-4.85, orange squares), or 1 mM EDTA (S-score:-8.07, red diamonds).

Source plates were pinned onto condition plates containing chemical stressors or subjected to environmental stresses after pinning, with a total of 3,026 mutants screened across 104 conditions in this way. After plate incubation, endpoint images were captured and processed, with colony size used as a proxy for fitness. Colony images were quantified using Iris software (36), which assigns a numerical colony size to each mutant in each condition. ChemGAPP software was then used to perform data normalisation, quality control, and calculation of S-scores (fitness scores) for each mutant–condition pair (37). S-scores capture the statistical difference between a mutant’s colony size in a specific condition and its median colony size across all conditions, enabling meaningful fitness comparisons between mutants that may differ in baseline growth (40).

Finally, the processed data can be explored using ChemGenXplore, a user-friendly interface that allows the scientific community to browse and visualise chemical genomics profiles for the *V. cholerae* C6706 library (available at https://chemgenxplore.kaust.edu.sa/) (32). ChemGenXplore enables users to visualise phenotypic profiles, assess gene-gene and condition-condition correlations, perform GO and KEGG enrichment analysis, and generate customisable, interactive heatmaps.

To maximise the number of fitness phenotypes detected, including those from genes with narrow stress responses, we designed the screen to include a broad set of diverse conditions targeting a wide range of cellular processes. In total, we obtained phenotypic profiles for 3,026 *V. cholerae* C6706 mutants tested across 104 conditions, corresponding to 73 unique conditions, for which some were tested at multiple concentrations (Figure 1B). These conditions included antimicrobial agents and other drugs used in human treatment that target diverse cellular processes in bacterial cells, such as cell envelope integrity, transcription, translation, and metabolic cofactor production (Figure 1B, bar chart expansion). In addition, we used alternative carbon sources, spent media from a range of bacteria (including *Vibrio cholerae*, *Acinetobacter baumannii*, and *Pseudomonas aeruginosa*), and environmental stresses such as altered growth temperatures and UV exposure. In general, we aimed to apply sub-inhibitory concentrations which reduce wild type growth to ≤ 50% as this limits the emergence of suppressor mutants.

### Screening dataset quality control

To assess the reliability of the raw screening data, we evaluated replicate reproducibility by calculating pairwise Pearson correlation coefficients between replicates. Replicates showed strong correlation, with an average coefficient of 0.746 (95% confidence interval: 0.745– 0.747), comparable to that reported previously for a similar screen in *E. coli* K-12 (21).

Next, we analysed the distribution of S-scores (fitness scores), which quantify how strongly a mutant’s growth in a given condition deviates from its growth across all other conditions in the screen. Under our null hypothesis, the *V. cholerae* C6706 mutant library contains only mutants inactivating non-essential genes, and these mutants are expected to respond significantly only in a small number of stress conditions where the inactivated gene contributes to fitness. Consistent with this expectation, S-scores were normally distributed (Figure S1), with most mutants showing few significant phenotypes in specific conditions, a pattern similar to that previously reported for *E. coli* K-12 (21). Whilst S-scores assign a relative fitness value to the same mutant across different conditions, they are also useful for comparing the growth of different mutants within a single condition. For example, the S-score distribution for pH 6 (Figure 1D) shows that mutants in *vqmA* (VCA1078) and *asd2* (VC2107) genes, which have negative S-scores, are visibly less fit than a mutant in *narP* (VCA0674), which has a positive fitness score.

To test whether S-scores capture relative fitness in liquid culture, we repeated selected conditions for the Δ*fadB* (VC1513) mutant, as this mutant exhibited fitness defects spanning mild to severe. The resulting growth curves show that more negative S-scores are associated with increasingly pronounced growth defects in liquid media. Overall, these tests give confidence that the generated S-scores can be used broadly as a proxy for mutant fitness in the respective conditions.

### Chemical genomic profiling of Vibrio cholerae C6706 library yields multiple robust phenotypes

After completing quality control on the chemical genomics dataset, we next asked what proportion of the mutant library exhibits significant phenotypes. S-scores provide relative fitness measurements for a mutant across all conditions, but the scores do not automatically indicate whether this is biologically relevant in a specific stress. To address this, we controlled the false discovery rate (FDR) at 5% and defined significant mutant-condition phenotypes as those with adjusted p-values below this threshold. We selected this 5% FDR cut-off based on previous chemical genomics studies in which a similar threshold was successfully applied (21,36). Remapping of library transposon insertion sites to the updated annotation resulted in a subset of mutants being reassigned to intergenic regions. To identify gene-specific phenotypes, these mutants were excluded from subsequent analysis. This resulted in a dataset for mutants in 2,748 non-essential genes, of which 2,589 (94.2%) were annotated, and 159 (5.8%) were unannotated, being listed as hypothetical or putative proteins.

Using a 5% FDR threshold, we detected 5,285 significant mutant–condition phenotypes, representing 1.76% of all 300,722 quality-controlled S-scores, with some mutants contributing multiple significant phenotypes (Figure 2A, blue line). Of these significant phenotypes, 2,616 (49.5%) were negative, indicating reduced growth under the tested condition, whereas 2,669 (50.5%) were positive, suggesting enhanced growth. We then examined how many significant phenotypes each mutant displayed (Figure 2A, green line). Most mutants with at least one phenotype did so in only one or two conditions, with 991 mutants (65.3% of all gene mutants exhibiting a phenotype) falling into this category. This is consistent with the idea that most genes contribute to fitness in a limited subset of environments, as seen in *Saccharomyces cerevisiae*, where 80% of deletion mutants have no effect in rich media but reveal fitness defects only when assayed across many diverse conditions (49).

**Figure 2.**
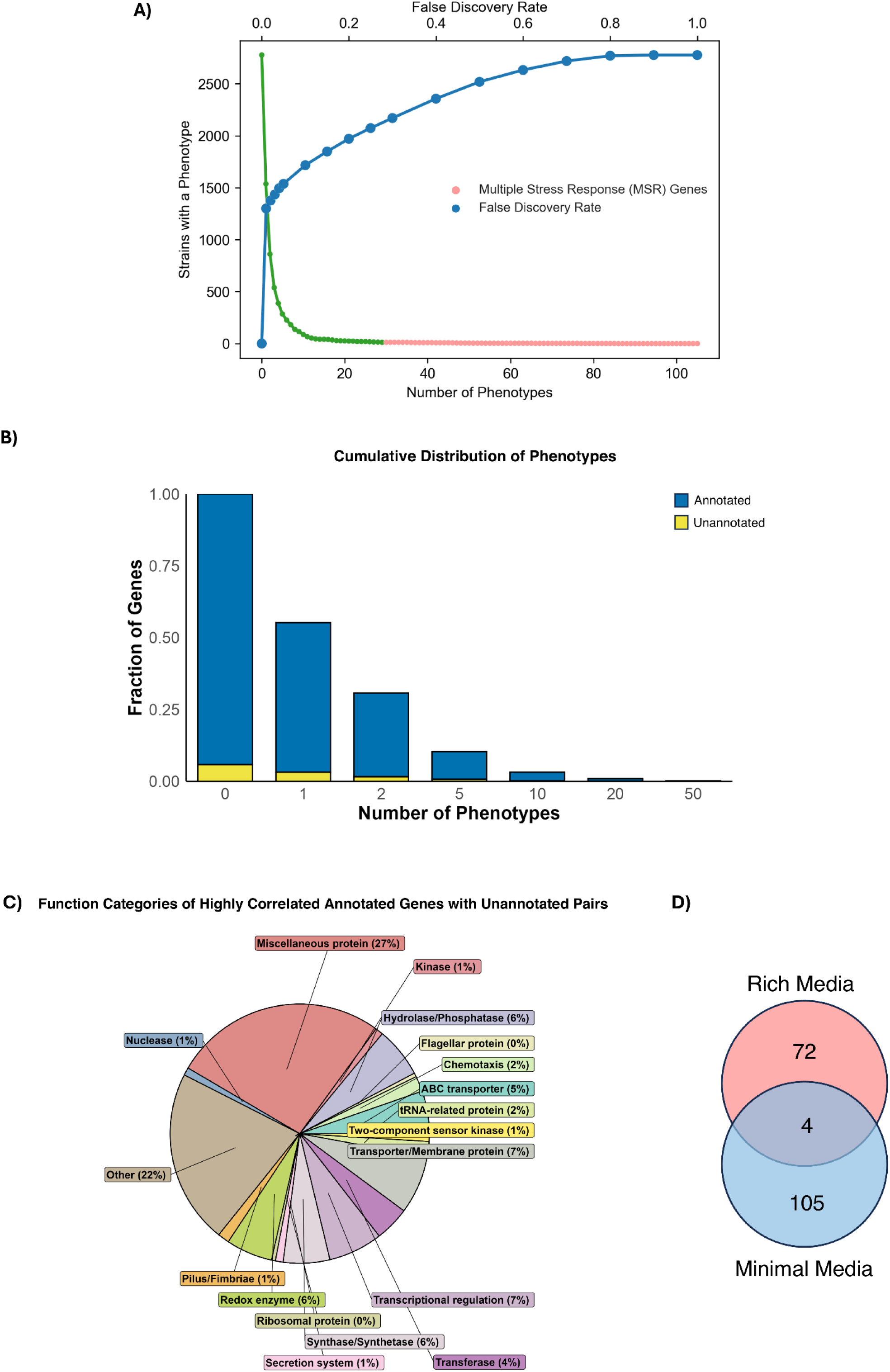
Identification and classification of phenotypes identified in screening dataset. **A)** As the False Discovery Rate (FDR) threshold is raised, the number of mutants with phenotypes increases (blue line). At an FDR cut-off at 5%, most mutants display one or two phenotypes, with smaller numbers having more than two phenotypes (green line). At an FDR of 5%, 55.2% of mutants were assigned at least one phenotype. Mutants with more than 30 phenotypes (pink line) are termed Multiple Stress Response Genes (MSRs). B) The histogram shows the cumulative distribution of significant phenotypes in annotated and unannotated genes, binned by the number of phenotypes identified. C) Pie chart showing functional categories of annotated genes highly correlated with unannotated genes. Annotated– unannotated gene pairs with strong phenotype similarity (r ≥ 0.7) are shown and are assigned only one function. D) Venn diagram showing numbers of conditionally essential genes required for growth in both rich and poor media conditions.

In contrast, a smaller subset of mutants showed significant phenotypes across multiple stress conditions (Figure 2A, pink line). Mutants with phenotypes in more than 30 conditions were classified as multiple stress-responsive (MSR) genes (Figure 2A, pink line). In total, we identified 12 MSR genes (Table S3), of which two — VC0025 and VC0115 — remain functionally uncharacterised in *V. cholerae*, VC0115 encodes a protein containing a domain of unknown function. Annotated MSR genes encompassed diverse functional categories, including genes associated with oxidative stress (e.g. *sodB*, encoding a superoxide dismutase), metabolism (e.g *fadE*, encoding an acyl-co-enzyme A dehydrogenase), and regulation of transcription and translation (e.g. VCA1078 encoding VqmA, a transcription factor involved in quorum-sensing (50), and VC1407, predicted to encode an RNA helicase) (Figure S2A).

### Identification of phenotypes for unannotated V. cholerae genes

A key objective of this study was to find functions for uncharacterised genes in *V. cholerae* through phenotypic profiling. We next assessed how the 5,285 significant fitness phenotypes were distributed between annotated and unannotated genes (Figure 2B). In total, we identified 1,430 annotated genes and 88 unannotated genes associated with phenotypes. A similar proportion of unannotated and annotated genes showed phenotypes: 55.3% of all tested unannotated gene mutants and 55.2% of annotated genes had at least one significant phenotype. This was somewhat unexpected, as genes contributing to fitness in defined conditions are more likely to have already been functionally annotated and experimentally confirmed. Additionally, unannotated genes tended to exhibit a similar number of significant phenotypes assigned per mutant compared to annotated genes (Figure S2B).

One approach to assigning gene function is to identify annotated genes with similar phenotypic profile. We therefore assessed the distribution of gene pairs with phenotypic profiles that correlated highly with each other. We found that a substantial fraction of unannotated– annotated gene pairs exhibited moderate to strong positive phenotypic correlations, with 10.5% of such pairs showing correlation coefficients of r ≥ 0.4 (Figure S2C). Figure 2C shows the functions of the annotated genes within highly correlated (r ≥ 0.7) gene pairs. We observed that phenotypes for many unannotated genes correlated with annotated genes involved in a wide range of cellular processes such as metabolism (including proteins with redox, synthetase and transferase activity), transport (ABC-transporters), envelope barrier function (membrane proteins and surface structures such as pilus-and fimbriae-associated genes), DNA metabolism (nucleases) and gene regulation (two-component sensor kinases and transcriptional regulators). Thus, our screen is effective for finding phenotypes for a wide variety of functional categories. Researchers wishing to find genes with strong phenotypic correlation to their gene of interest can do so using the correlation analysis function of ChemGenXplore (32).

In some cases, genes become essential for cell viability in a given condition as gene disruption results in extremely poor growth, reflected in highly negative S-scores. For example, in a minimal media condition, disruption of a conditionally essential gene may render a strain auxotrophic. We identified a total of 192 conditionally essential genes in our dataset (Figure 2D), which were required for growth in both nutrient-restricted and rich-media conditions with a small subset included in both categories (a list of conditionally essential genes is available in Table S4). Hence, the phenotypes we identified were not restricted to either rich-or restrictive-media conditions.

### Mining the dataset to find biology for individual mutants

After confirming that the dataset can detect robust phenotypes for unannotated genes, we next decided to focus on an example to demonstrate how phenotypes can be used to suggest gene function. For this purpose, we selected VCA0734, which is encoded on the smaller second chromosome of *V. cholerae* C6706. In our screening dataset, the ΔVCA0734 mutant displayed a negative fitness phenotype in multiple envelope stress conditions including the large glycopeptide antibiotics vancomycin and bleomycin, isopropanol, low pH conditions, and the disinfectant benzalkonium chloride (Figure 3A). To ensure that our screening results were not an artefact, we carried out growth curves for the ΔVCA0734 transposon mutant in LB medium with or without 5% isopropanol, which confirmed the phenotype (Figure 3B). According to AlphaFold3 predictions (51), the VCA0734 gene appears to encode a 10-stranded β-barrel protein (Figure 3C), and BLAST searches indicate that it is phylogenetically restricted to the family *Vibrionaceae*. The function of the VCA0734 gene product remains unclear, although previous studies have linked its expression to motility through the histidine kinase FlrB (52), norspermidine signalling (53), and the regulatory region is targeted by the alternative σ-factor RpoN (54). VCA0734 also appears to be subject to negative regulation by the iron-responsive regulator Fur, although this regulation occurs independently of iron (55). Despite our text mining efforts, the current literature provides no clear clues to the cellular function of VCA0734.

**Figure 3.**
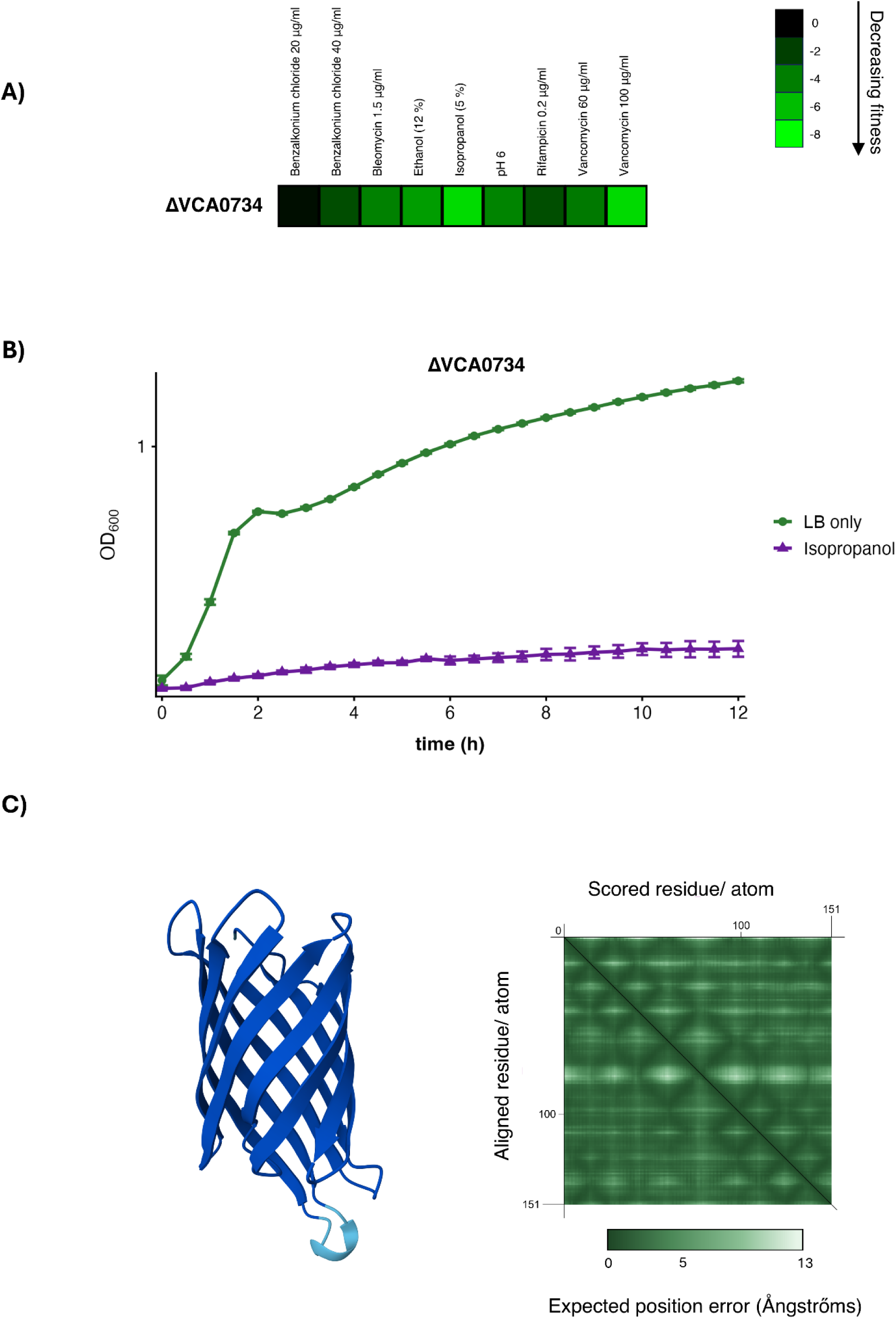
VCA0734 is a predicted outer membrane protein required for fitness in envelope stresses. A) Heat map showing phenotypic profile of ΔVCA0734 mutant in selected conditions. Higher intensities of green indicate decreasing fitness. B) Growth curve for a *V. cholerae* C6706 ΔVCA0734 transposon mutant grown in LB media (green circles) or LB media supplemented with 5% isopropanol (purple triangles). C) Alphafold (51) prediction for VCA0734 and PAE chart showing degree of confidence per amino acid position. pLDDTs are colour-coded on the model (dark blue > 90 and cyan for 70-90), the pTM for the prediction is 0.9. The input sequence has been clipped after the predicted signal peptide (56). PAE chart was generated using PAE viewer (100).

Using SignalP (56), the VCA0734 protein is predicted to carry a Sec signal peptide (likelihood 0.9953), indicating that VCA0734 is exported to the periplasm. The observed sensitivity of a ΔVCA0734 mutant to envelope stressors – especially large glycopeptide antibiotics, which typically cannot penetrate the Gram-negative outer membrane (57) – suggests a defect in envelope integrity. Taken together with the AlphaFold model, we speculate that VCA0734 likely contributes to the barrier function of the outer membrane. Experimental validation for this hypothesis is still needed, though we hope this example illustrates how scientists can approach and use this chemical genomics dataset as a resource.

### Network analysis of chemical genomics dataset identifies functional clusters

Phenotypically related genes can be clustered together, revealing common gene sets that are important for cellular pathways. Thus, this approach provides a broader view of gene function within the cell. Similarly, clustering stress conditions can reveal linked gene groups, providing a broader view of cellular responses. Such network-based analyses can be performed in two or three dimensions, offering complementary perspectives on how genes and conditions are functionally related. Examples of each type of analysis are shown in Figure 4. To increase stringency, only mutants with transposon insertions located within the central 50% of a gene coding sequence (i.e. between 25–75% of gene length) were considered, thereby maximising the likelihood of loss-of-function effects.

**Figure 4.**
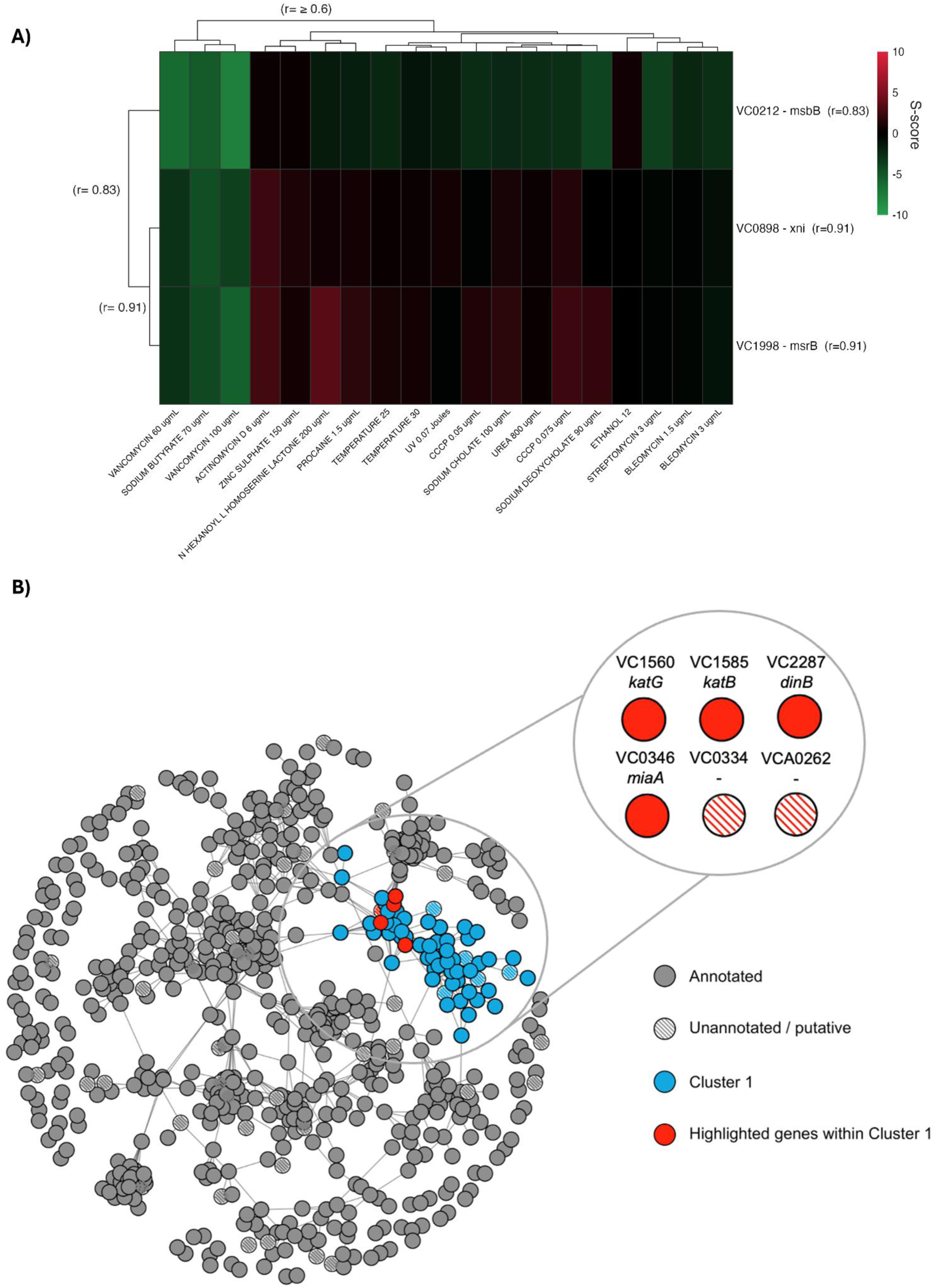
Network analysis of phenotypically related genes in *V. cholerae* using two-and three-dimensional clustering. A) Two-dimensional clustering of a subset of phenotypically related genes in highly correlated conditions (r ≥ 0.6). Correlation coefficient values in parenthesis show the correlation of the gene to its nearest neighbours within the cluster. B) A gene–gene correlation network was constructed from chemical genomics phenotypes using Pearson correlation (r ≥ 0.6). Nodes represent genes and edges indicate strong phenotype similarity. Annotated genes are shown as filled nodes, and unannotated genes are cross-hatched. Network communities were identified using the Louvain community detection algorithm, and the largest community (cluster 1) is highlighted in blue. Selected genes from this cluster shown in the expansion are coloured red.

Figure 4A shows an example of genes whose phenotypes cluster in two dimensions across a subset of conditions. These conditions include cell-envelope stresses caused by antibiotics such as vancomycin (58), detergents such as sodium deoxycholate, and the proton motive force uncoupler CCCP (carbonyl cyanide m-chlorophenyl hydrazone) (59). Other conditions include DNA-damaging agents, including UV irradiation (60) and the glycopeptide antibiotic bleomycin (61). Sodium butyrate is also included in this cluster and has been reported to interfere with virulence regulation in *V. cholerae* via the ToxT protein (62), as well as to affect membrane function and proton motive force maintenance (63).

In the heatmap, mutants in three genes form a cluster with similar phenotypic patterns across the conditions shown. The *msrB* and *xni* genes cluster closely together, which is consistent with a link between oxidative stress and DNA repair functions: *msrB* encodes a methionine sulfoxide reductase involved in protein quality control under oxidative stress (64), and *xni* which encodes a flap endonuclease. Flap endonucleases have 5′-endonuclease activity, participate in resolution of RNA-DNA hybrids, and are implicated in DNA replication and repair (65,66). These genes cluster with *msbB*, which encodes a lipid A acyltransferase required for resistance of the El Tor biotype of *V. cholerae* to polymyxin B and a number of antimicrobial peptides, a property not shared by the classical biotype (67). In *V. fischeri*, MsbB is reported to promote resistance to detergents, antimicrobial peptides, and antibiotics (68). Overall, we speculate that this cluster reflects coordinated stress-response programs that may connect cell-envelope maintenance with DNA and protein repair.

Alternatively, three-dimensional clustering enables a more holistic view of phenotypic interactions. An example of this is displayed in Figure 4B, together with an expanded subset of clustered genes whose mutants also show similar phenotypic profiles. This subset contains multiple stress-response genes involved in responding to reactive oxygen stress and DNA damage. These include *katG* (VC1560) and *katB* (VC1585), two genes encoding a catalase-peroxidase and catalase enzyme respectively (69), and *dinB*, which encodes DNA polymerase IV, an error-prone translesion polymerase that plays an important role in the SOS response (70). The *miaA* gene also falls in this cluster and encodes a tRNA-modifying enzyme that catalyses the first step required for the ms^2^i^6^ modification at position A-37 on tRNAs that recognise UNN codons (71). This modification prevents translation frame-shifting, and *miaA* mutants in *Salmonella* enterica serovar Typhimurium exhibit growth defects and altered translation rates, whereas an *E. coli miaA* mutant displays recombination-dependent mutator phenotype (72–74). Hence, *miaA* is associated with genome stability and is also linked to general stress responses through its requirement for efficient translation of *rpoS* and *hfq* mRNA, which are both involved in stress adaptation (75–77). Two unannotated genes, VC0334 and VCA0262, also cluster closely with these stress-response genes. VC0334 shows homology to acetyltransferases, and VCA0262 encodes a protein containing a domain of unknown function (DUF2946). Taken together, this clustering approach can highlight phenotypically relevant unannotated genes that may provide leads for future functional studies.

### ChemGenXplore: an app enabling exploration of the dataset

To facilitate the use of our dataset as a resource, the data is browsable within ChemGenXplore (32), and is available at https://chemgenxplore.kaust.edu.sa/. As an overview, ChemGenXplore enables visualisation of phenotypes for genes of interest, filtering of the data by FDR thresholds and transposon insertion position, identification of gene-gene or condition-condition correlations, and functional classification using GO terms or KEGG pathways that may be enriched in a particular set of conditions (Figure 5, Figure S3). Finally, the software provides a wide variety of options for visualising data in formats suitable for publication.

**Figure. 5.**
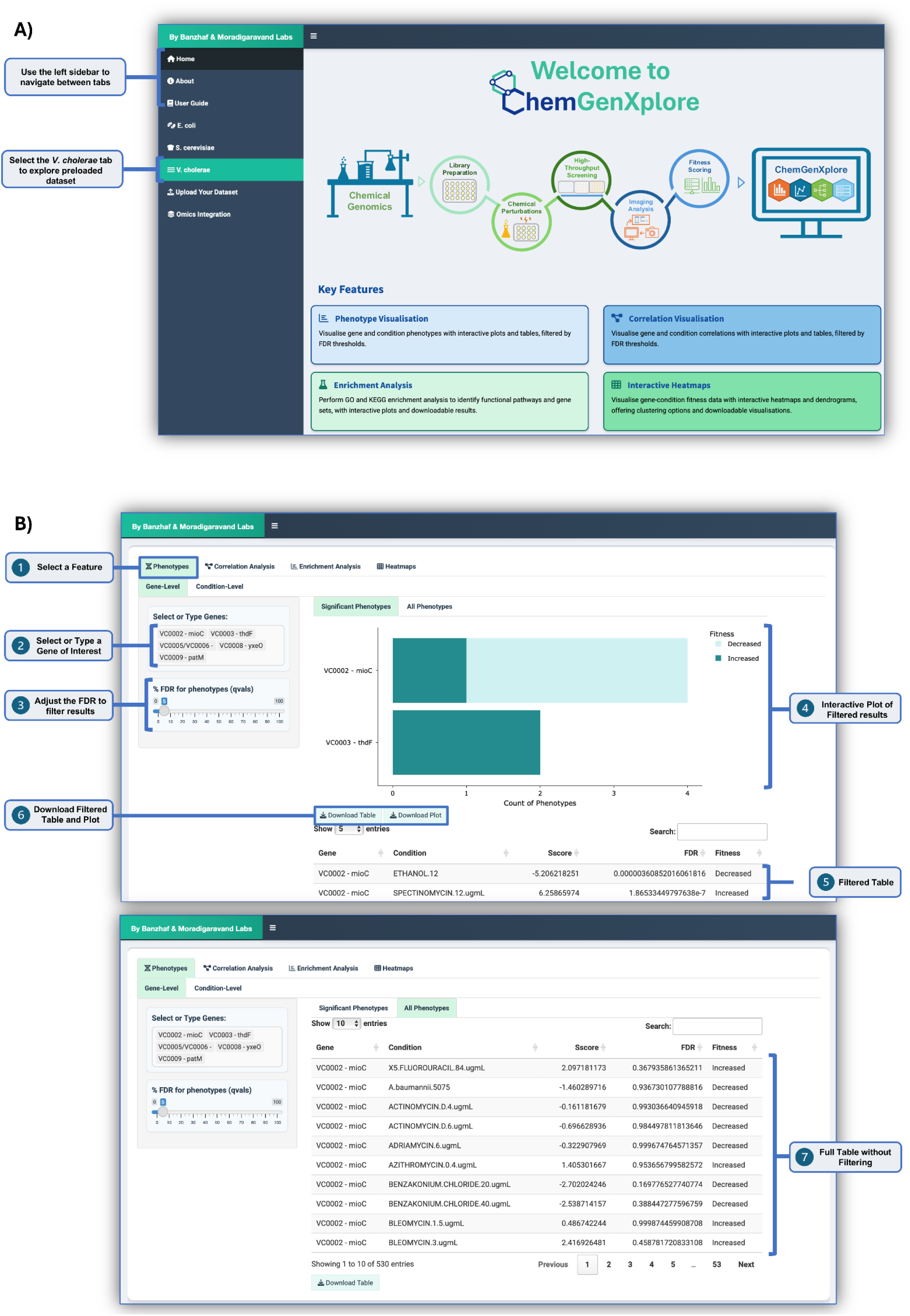
ChemGenXplore as a tool to explore chemical genomics datasets. A) Users can either browse existing datasets or upload their own. B) ChemGenXplore enables exploration of strain phenotypes or condition, in addition to a variety of analysis tools (further illustrated in Figure S3).

## DISCUSSION

For genomic sequencing data to translate into biological insight, the functions of individual genes need to be understood. This is particularly important for bacterial pathogens such as *Vibrio cholerae*, which occupy environmental reservoirs outside the host and rely on diverse mechanisms to persist and transition between niches. *V. cholerae* also serves as a key model for studying quorum sensing, pathogenesis, and gene regulation (78–82). However, many genes involved in these processes are either hypothetical or lack experimental validation of their function, and this knowledge gap limits understanding of the biology of this pathogen.

We applied a chemical genomics pipeline (20) to systematically profile fitness phenotypes in a single-gene mutant library of *V. cholerae* strain C6706 across 104 conditions. The screen identified 5,285 significant gene-condition associations, substantially expanding the available functional dataset for *V. cholerae*. Using a 5% false discovery rate, we identified phenotypes for 55.2% of mutants tested, a similar proportion to that reported by Nichols *et al*. in *Escherichia coli* using the same threshold (21). Notably, 88 out of 159 (55.3%) tested unannotated gene mutants displayed phenotypes— a relatively high proportion given that genes with easily detected phenotypes are more likely to have been previously characterised and annotated. The distribution of phenotypes for unannotated genes was broadly similar to that of annotated genes, contrasting with *E. coli*, where unannotated genes tend to show fewer phenotypes (21).

Whereas the *E. coli* screen identified 102 genes that were conditionally essential in rich media, and 116 in minimal media, with 22 genes overlapping these two categories (21), we found that *V. cholerae* harbours comparable conditionally essential genes in rich media (75 genes) and minimal media (121 genes), but with substantially less overlap (4 genes). This difference may reflect organism-specific metabolism, physiology and niche adaptation between *V. cholerae* and *E. coli*. In particular, *V. cholerae* is adapted to fluctuating aquatic and host-associated environments and can utilise substrates such as N-acetylglucosamine and citrate (83–86), whereas *E. coli* has broader metabolic flexibility (87–89).

The high proportion of phenotypes for unannotated genes in our dataset may further reflect the importance of niche-and condition-specific functions in the lifestyle of *V. cholerae*. Although largely clonal, pandemic *V. cholerae* strains demonstrate significant genome plasticity, acquiring mobile genetic elements that enable niche survival (90–93). For example, the VSP-II (Vibrio Seventh Pandemic) island contains genes responsive to zinc starvation, which are not induced under standard laboratory conditions (94). Such genes may lack clear phenotypes in traditional molecular biology assays or homologs in model organisms and therefore remain uncharacterised. Previous large-scale fitness screens have provided valuable insight into *V. cholerae* biology, but were typically restricted to defined environments (95), or focused on specific phenotypic outputs (96,97) rather than systematically surveying fitness across a broad panel of diverse conditions. Thus, this dataset illustrates the utility of systematic chemical genomics screens for prioritising genes for follow-up study that are potentially relevant to pathogen biology.

However, this study has several limitations. Transposon mutagenesis can produce unintended polar effects on downstream genes within operons (31), and the resulting libraries cannot capture the roles of essential genes, as stable mutants cannot be generated for these loci. We therefore recommend that users confirm the position of transposon insertions (available in Table S1) when interpreting gene-specific phenotypes. In addition, the high-throughput format of the chemical genomics pipeline constrains the range of conditions that can be tested, making it challenging to reproduce complex environments that mimic natural or host-associated niches of *V. cholerae*. For example, pandemic *V. cholerae* strains encode multiple type VI secretion systems and natural transformation machinery for uptake of prey DNA, which confer competitive advantages only in specific niches (8,93,98,99), and such functions are difficult to assess in this format. Hence, our dataset reflects the 104 conditions tested rather than the full ecological breadth of the organism.

Despite these limitations, this dataset should be useful to the community as a searchable resource for identifying phenotypes in both annotated and unannotated genes, thereby guiding hypothesis-driven research. The ChemGenXplore platform is designed to make the data readily accessible, enabling researchers to quickly explore the screen, or upload and analyse their own datasets, without requiring prior bioinformatics expertise.

## ACKNOWLEDGEMENTS

We thank Kai Papenfort for donating a copy of the *V. cholerae* C6706 single-gene mutant library.

## SUPPLEMENTARY TABLES

**Table S1: Strains used in this study.**

**Table S2: Screening conditions.**

**Table S3: MSR genes.**

**Table S4: Conditionally essential genes.**

