## Supplementary Figures for "Systematic chemical genomic screening of *Vibrio cholerae* maps gene-phenotype relationships"

**Figure S1**

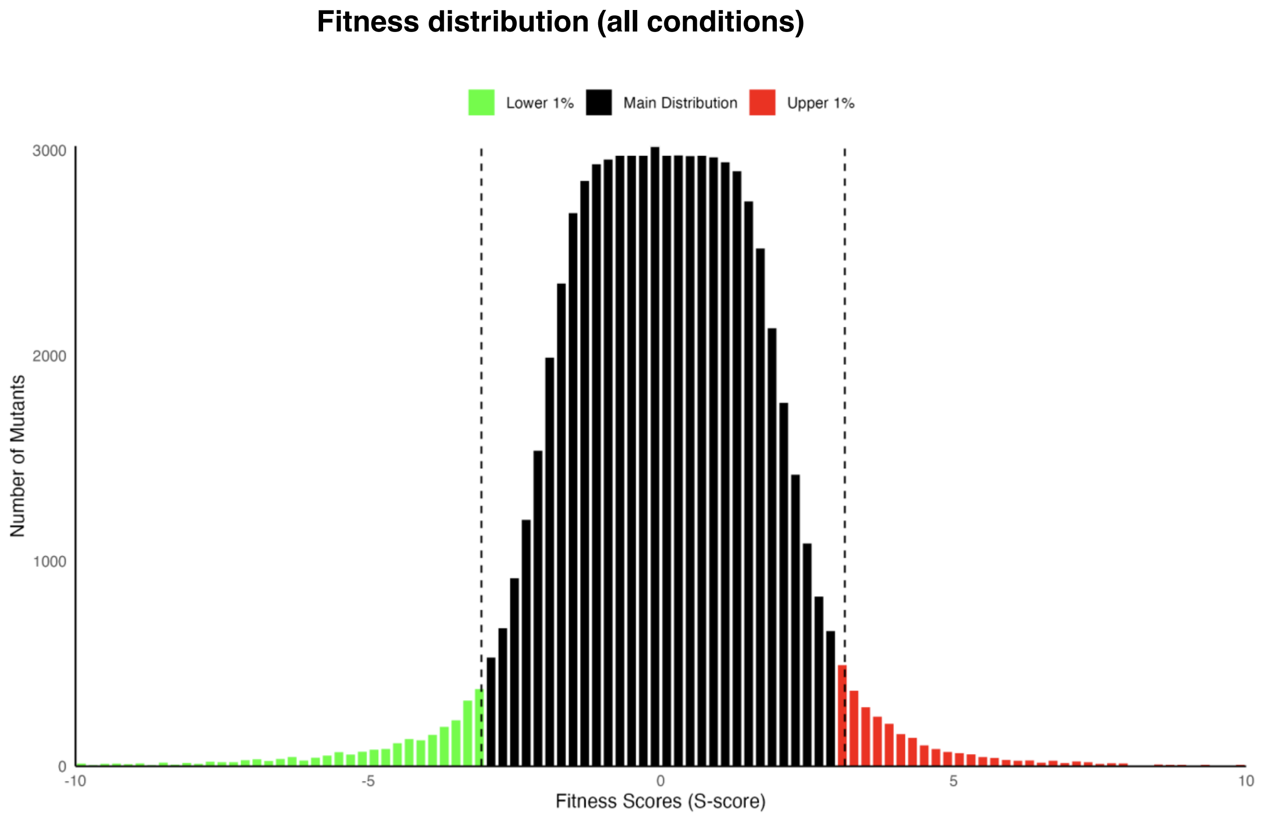
Fitness distribution (all conditions)
Lower 1%
Main Distribution
Upper 1%
3000
2000
1000
0
Number of Mutants
-10
-5
0
5
10
Fitness Scores (S-score)

**Figure S2**

**Function Groups of MSR Genes (≥30 significant phenotypes; FDR ≤ 0.05)**

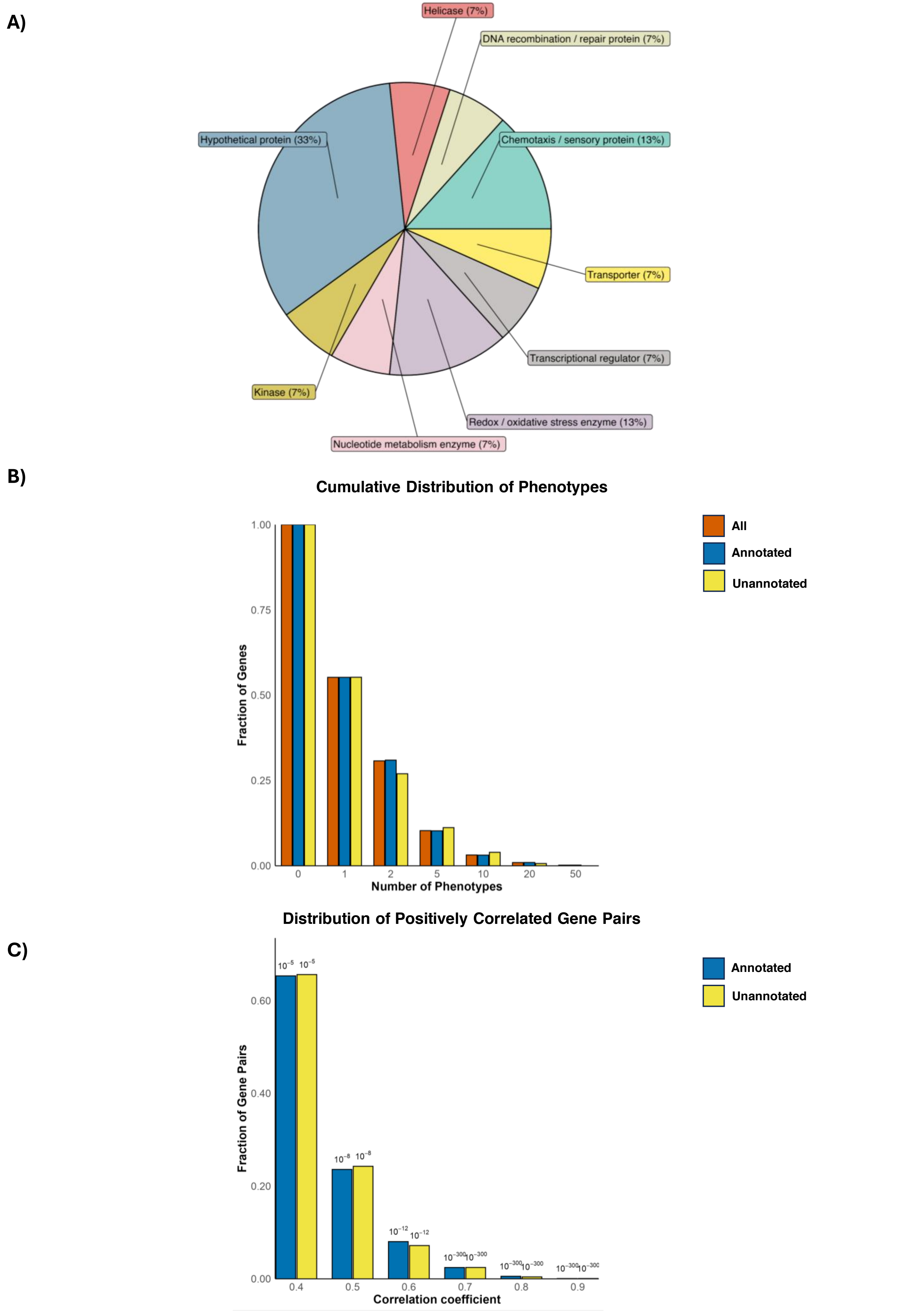

**Figure S3**

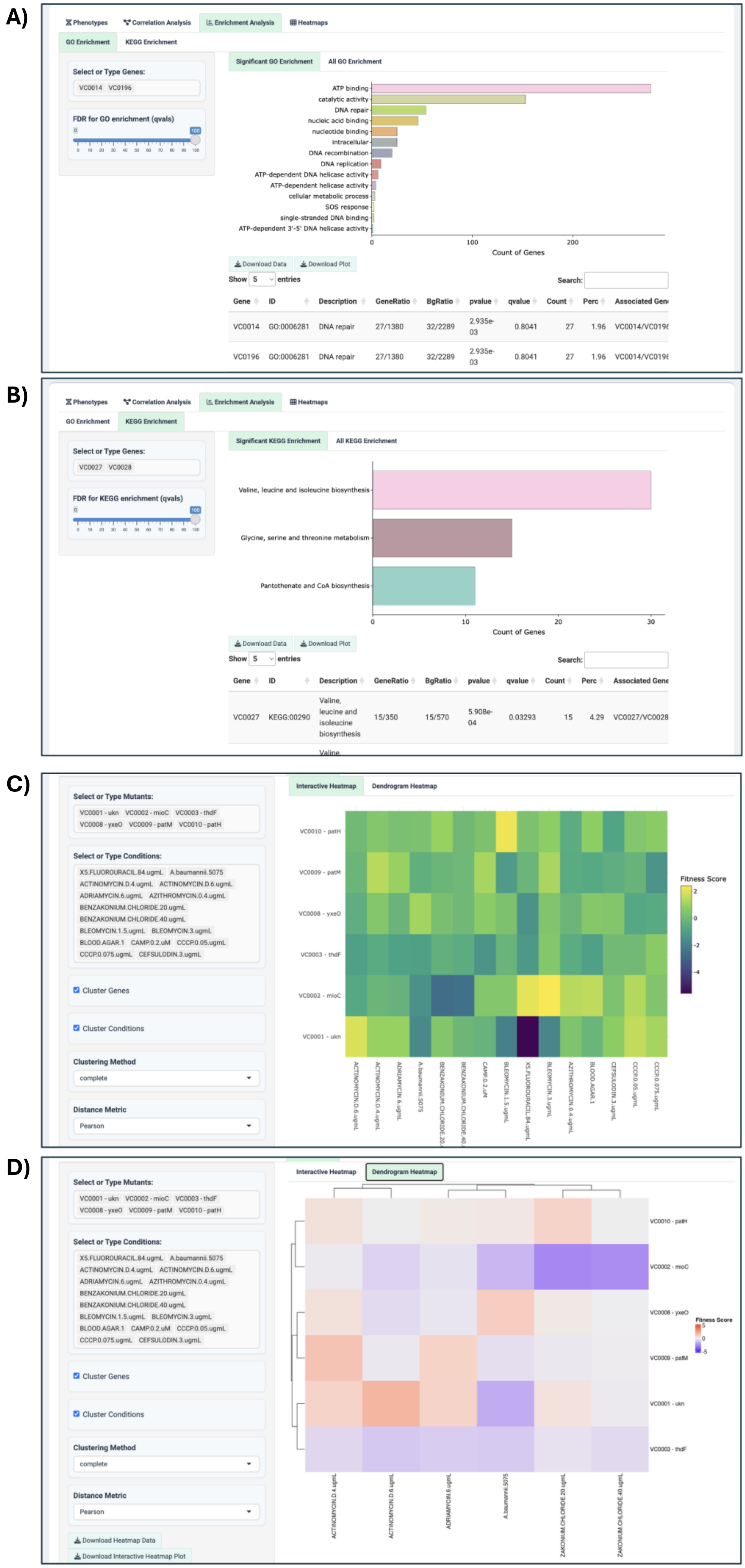

A)
Phenotypes
Correlation Analysis
Enrichment Analysis
Heatmaps
GO Enrichment
KEGG Enrichment
Significant GO Enrichment
All GO Enrichment
Select or Type Genes:
VC0014 VC0196
FDR for GO enrichment (qvals)
ATP binding
catalytic activity
DNA repair
nucleic acid binding
nucleotide binding
intracellular
DNA recombination
DNA replication
ATP-dependent DNA helicase activity
ATP-dependent helicase activity
cellular metabolic process
SOS response
single-stranded DNA binding
ATP-dependent 3'-5' DNA helicase activity
Count of Genes
Download Data
Download Plot
Show 5 entries
Search:
Gene ID Description GeneRatio BgRatio pvalue qvalue Count Perc Associated Gen
VC0014 GO:0006281 DNA repair 27/1380 32/2289 2.935e-03 0.8041 27 1.96 VC0014/VC019
VC0196 GO:0006281 DNA repair 27/1380 32/2289 2.935e-03 0.8041 27 1.96 VC0014/VC019
B)
Significant KEGG Enrichment
All KEGG Enrichment
VC0027 VC0028
FDR for KEGG enrichment (qvals)
Valine, leucine and isoleucine biosynthesis
Glycine, serine and threonine metabolism
Pantothenate and CoA biosynthesis
Count of Genes
VC0027 KEGG:00290 Valine, leucine and isoleucine biosynthesis 15/350 15/570 5.908e-04 0.03293 15 4.29 VC0027/VC0028
C)
Select or Type Mutants:
VC0001 - ukn VC0002 - mioC VC0003 - thdF VC0008 - yxeO VC0009 - patM VC0010 - patH
Select or Type Conditions:
X5.FLUOROURACIL.84.ugmL A.baumannii.5075 ACTINOMYCIN.D.4.ugmL ACTINOMYCIN.D.6.ugmL ADRIAMYCIN.6.ugmL AZITHROMYCIN.0.4.ugmL BENZAKONIUM.CHLORIDE.20.ugmL BENZAKONIUM.CHLORIDE.40.ugmL BLEOMYCIN.1.5.ugmL BLEOMYCIN.3.ugmL BLOOD.AGAR.1 CAMP.0.2.uM CCCP.0.05.ugmL CCCP.0.075.ugmL CEFSULODIN.3.ugmL
Cluster Genes
Cluster Conditions
Clustering Method
complete
Distance Metric
Pearson
Interactive Heatmap
Dendrogram Heatmap
Fitness Score
D)
Download Heatmap Data
Download Interactive Heatmap Plot

### SUPPLEMENTARY FIGURE LEGENDS

**Figure S1. Histogram showing distribution of S-scores for across all mutants across all screening conditions.** The lowest 5% of S-scores (indicating fitness) are shown in green, the top highest 5% of positive S-scores are shown in red.

**Figure S2.**

A) **Functional categories of MSR genes identified in *V. cholerae* chemical genomics screen.** Each gene is assigned one function.

B) **Cumulative distribution of phenotypes expressed as a fraction of each gene class.** Genes are binned according to the number of phenotypes identified and have at least the number of phenotypes of the x-axis category.

C) **Distribution of positive phenotypic correlations across genes.** The histogram shows the fraction of gene pairs correlated with annotated genes, binned by correlation coefficient. Numbers above bars indicate the median correlation p-value for each bin.

**Fig S3. Additional functionality of ChemGenXplore.** In addition to exploring strain phenotypes, ChemGenXplore also enables users to carry out enrichment analysis using GO (A) or KEGG terms (B), generation of interactive heatmaps (C) and dendrograms to highlight clustering of conditions or phenotypically related genes (D).
